# Extracellular K^+^ modulates the pore conformations of Cys-loop receptor anion channels

**DOI:** 10.1101/2025.06.15.658987

**Authors:** Takushi Shimomura, Yoshihiro Kubo, Minoru Saitoe, Yoshinori Suzuki

## Abstract

K^+^ is an essential cation for life, but no eukaryotic membrane protein with a modulatory site for extracellular K^+^ has been discovered. Here, we report that a Cys-loop receptor, CG12344/DmAlka, expressed in the *Drosophila* nervous system, is selectively modulated by physiological concentration of extracellular K^+^. Structural prediction, electrophysiology and phylogenetic analysis of DmAlka revealed the extracellular K^+^ binding site that mimics the hydrated chemical environment for K^+^, as observed in K^+^ channel pore. Furthermore, we found that K^+^ binding induces a previously unrecognized “mode-switching,” altering properties ranging from ligand sensitivity to ion selectivity. Notably, a human glycine receptor variant also exhibited similar mechanisms. Our study reveals a novel regulatory mechanism of Cys-loop receptors that directly links the extracellular K^+^ signaling to Cl^-^ conductance in animals.

## Introduction

Potassium ions (K^+^) are essential cations for all living organisms, playing a vital role in multiple cellular functions, such as pH homeostasis, osmoregulation, cofactors for enzymatic activities, and especially in generating membrane potential (*1–3*). In order to set the resting membrane potential appropriately, typical extracellular concentrations of K^+^ in mammals are regulated in the range of 3-5 mM, which underlies the membrane excitability of neuronal and muscular cells (*4*, *5*).

Changes in extracellular K^+^ concentrations are generally detected and maintained by K^+^-permeable membrane proteins, such as channels, transporters, and pumps, with enhancement of their K^+^ permeability depending on K^+^ itself or through the shift in membrane potential caused by K^+^ influx (*2*). Besides these membrane proteins that utilize K^+^ as substrates, only a few proteins are known to have molecular modules for responding selectively to extracellular K^+^ concentrations. In eukaryotes, plants have been known to have systems for detecting changes in soil K^+^ concentrations, although the molecular identities of their K^+^ sensors are unclear (*6–8*). In animals, to the best of our knowledge, no proteins have been reported to detect extracellular K^+^ directly and selectively as a regulatory signal, rather than as a substrate.

Here, we report that a *Drosophila* Cys-loop receptor-type of chloride channel, CG12344, is selectively regulated by physiological concentration of extracellular K^+^. Structure prediction combined with electrophysiological analysis revealed a structural determinant for its K^+^ selectivity. We also found that, with an unprecedented mechanism among Cys-loop receptors, K^+^ modulates the *Drosophila* channel and also a variant of the human glycine receptor. These findings suggest a novel signaling pathway connecting extracellular K^+^ to Cl^-^ conductance through the Cys-loop receptors.

## Results

### DmAlka is selectively modulated by the physiological concentration of extracellular K^+^

CG12344 was previously considered to be a homolog of glycine receptors (GlyRs) (*9*). However, recent studies have shown that it is not activated by glycine but is sensitive to alkaline pH, and therefore it has been named *alkaliphilie* (DmAlka) (*10*). We used the *Xenopus* oocyte expression system to characterize the molecular details of DmAlka. Consistent with the previous report using mammalian cells (*10*), DmAlka evoked Cl^-^ currents in a standard bath solution without glycine in *Xenopus* oocytes (fig. S1A, B). We newly found that DmAlka currents increase in accordance with the decrease in the extracellular K^+^ concentration (Fig. 1A). The dependence of extracellular K^+^ was observed in the physiologically relevant concentrations (Fig. 1B). Evaluation of the effects of different alkali-metal cations in 2 mM revealed a selectivity sequence similar to that of K^+^ channels (Rb^+^ ∼ K^+^ > Cs^+^ >> Na^+^ ∼ Li^+^) (*11*) (Fig. 1C, D and fig. S_1_C). Change in Na^+^ concentration hardly affected Cl^-^ currents, while K^+^ significantly changed it, demonstrating the K^+^ selective modulation of DmAlka under the physiological concentrations of extracellular Na^+^ and K^+^ (fig. S1D).

**Fig. 1.**
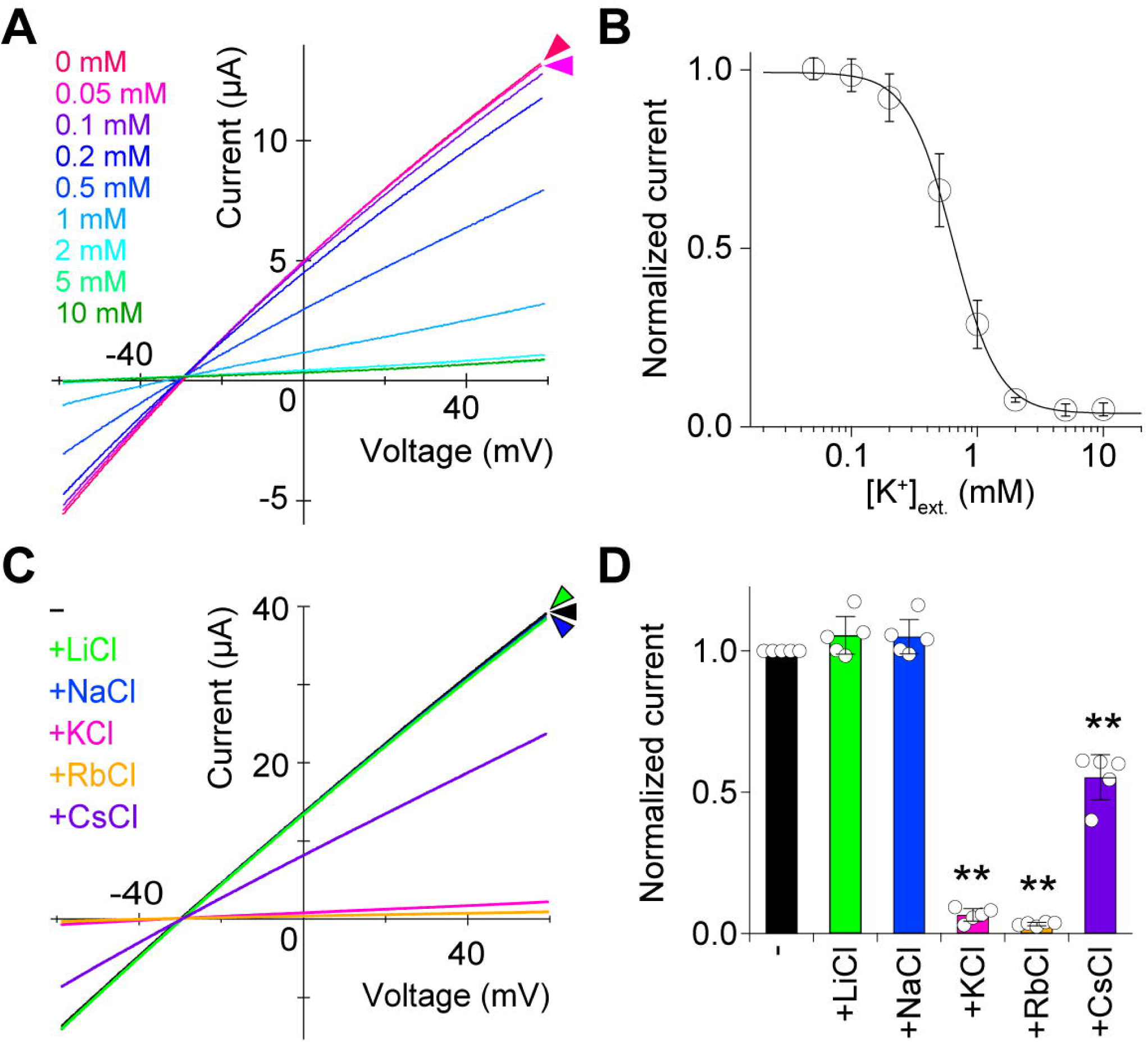
Extracellular K^+^ dependence of DmAlka. (**A**) Representative current traces of DmAlka evoked by ramp pulses from –60 to +60 mV under various concentrations of extracellular K^+^ (0–10 mM) in the standard bath solution ND96. The traces are color-coded gradually from magenta (low concentration) to green (high concentration). (**B**) A dose-response curve of the extracellular K^+^-induced reduction in DmAlka currents (IC_50_ = 0.645 ± 0.019 mM, n = 4). (**C**) Representative current traces recorded in the ND96-based various external solutions, in which 96 mM NaCl was replaced with NMDG-Cl, and each of 2 mM monovalent cations or no cations is included. The cation chlorides tested were LiCl (green), NaCl (blue), KCl (magenta), RbCl (orange), and CsCl (purple). (**D**) Plots of the current amplitude at +60 mV in the presence of each 2 mM monovalent cation, normalized to the value obtained in their absence, as shown in (C) (n = 5). Color codes correspond to those used in (C).

### An AlphaFold3 model of DmAlka predicts the K^+^ selective binding site

To gain insight into the structural basis of K^+^-dependent modulation in DmAlka, we used AlphaFold 3 (AF3) (*12*). AF3-based structural prediction of DmAlka in a K^+^-bound state, which has a confident reliability score (fig. S2A), revealed an overall structure of typical Cys-loop receptors, characterized by its pentamer stoichiometry and a conserved Cys-loop (Fig. 2A) (*13*, *14*). The predicted structure was composed of the extracellular domain (ECD), which receives neurotransmitters such as glycine for GlyRs, and the transmembrane domain (TMD) that forms the ion-permeable pore.

**Fig. 2.**
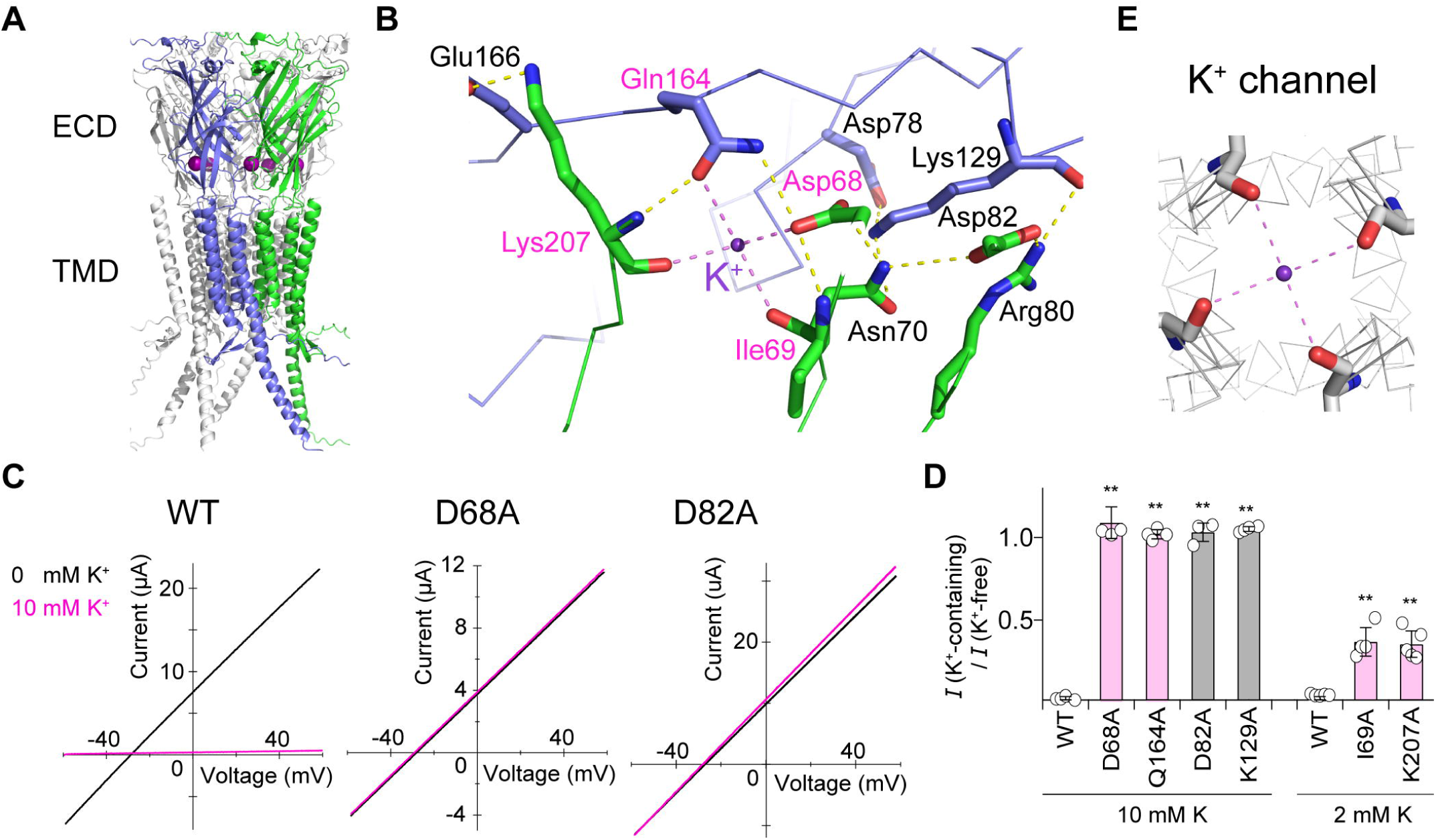
Structural basis of K^+^ selectivity in DmAlka predicted by AF3. (**A**) An AF3-predicted structure of DmAlka. K^+^ are represented as purple spheres. Two adjacent subunits are colored in navy blue and green. (**B**) A close-up view of the K^+^- binding site shown in (A). Side-chain and main-chain atoms involved in K^+^ binding are shown as stick models with main-chain traces visualized in ribbon model. Coordination of K^+^ with four oxygen atoms and the polar interactions surrounding the K^+^ binding are indicated by dashed lines colored in purple and yellow, respectively. (**C**) Representative current traces of DmAlka WT, D68A, and D82A mutants, recorded in the absence (black) or presence (magenta) of 10 mM extracellular K^+^. (**D**) K^+^-induced current inhibition in WT and various mutants. The bars colored in pale magenta indicates the mutations to the residues directly involved in K^+^ coordination (n = 3–5). (**E**) The selectivity filter in the crystal structure of the KcsA K^+^ channel (PDB: 1K4C). K^+^ is shown in purple. Interactions with four coordinating oxygen atoms of Tyr78, shown as stick model, are represented as purple dashed lines.

The K^+^ binding site is formed between three loops —Cys-loop, and C-terminus of β1 strand connecting to β1-β2 loop and β8-β9 loop (named according to the nomenclature of Cys-loop receptors) —in the ECDs (Fig. 2A, fig. S2B) (*15*). These loops formed an interface that connects the ECD to TMD. An extensive network of polar interactions, composed of hydrogen bonds and salt bridges, is formed surrounding the K^+^ binding site (Fig. 2B). K^+^ is directly coordinated to carbonyl or carboxyl oxygen atoms of Asp68, Ile69, Gln164, and Lys207. Mutations of the residues that directly form or indirectly stabilize the binding site through the polar network impaired K^+^ sensitivity, depending on the degree of their contribution at the site (Fig. 2C, D, fig. S2C). Phylogenetic analysis of DmAlka homologs revealed that the four key K^+^-coordinating residues, along with surrounding residues involved in the polar interactions, are well conserved across the arthropod phylum (fig. S3A-C). The AF3 models of these homologs showed the same K^+^ binding sites with high confidence as that of DmAlka (fig. S3D). These results validate the predicted DmAlka model and suggest that the K^+^-recognition mechanism in DmAlka is physiologically conserved within the arthropods.

The electrostatic surface potential of DmAlka revealed that, unlike typical Cl^-^ channels that have a positively charged potential close to the pore, the region connecting to the K^+^ binding site is negatively charged, presumably optimal for attracting cations (fig. S2D). A detailed inspection of the K^+^ binding site revealed that distances between K^+^ and its coordinating oxygen atoms are within 2.6-3.0 Å across all five K^+^ in a pentamer (2.79 ± 0.13 Å in average), which closely matches the coordination distance between K^+^ and water oxygen atoms (approximately 2.8 Å) (*16*). This is similar to the structure and mechanism of K^+^ selectivity in K^+^ channel pore, in which Na^+^ faces a higher energetic barrier to remove its coordinated water molecules and to enter the selectivity filter due to its less favorable coordination distance with oxygen atoms than K^+^ (*17*) (Fig. 2B, E). Taken together, these findings show that DmAlka has an efficient architecture for K^+^-dependent modulation, especially highlighted by the K^+^ binding site mimicking the hydrated chemical environment for K^+^.

### A K^+^-dependent mode-switching mechanism in DmAlka

In DmAlka expressed in *Xenopus* oocytes, moderate alkalization enhanced DmAlka currents as previously reported (*10*), and stronger alkalization resulted in the current increase, followed by the gradual decrease due to desensitization in the presence of K^+^ (Fig. 3A, fig. S4A). We newly found that this alkaline-dependence was impaired when the K^+^ concentration was reduced, making DmAlka more pH-independent, non-desensitizing, and constitutively open (Fig. 3A, B, fig. S4A). The K^+^ dependence of alkaline sensitivity was also confirmed using the D82A mutant, which has been shown to lack the K^+^ dependent inhibition (Fig. 2C, D). D82A exhibited constitutively active, alkaline-independent currents irrespective of the K^+^ presence (fig. S4B, C). Conversely, the M77R mutant, which was engineered to introduce a positive charge at the K^+^ binding site and therefore mimics a constitutively K^+^-bound state, retained alkaline sensitivity even in the absence of K^+^ (Fig. 3C, D, fig. S4D, E).

**Fig. 3.**
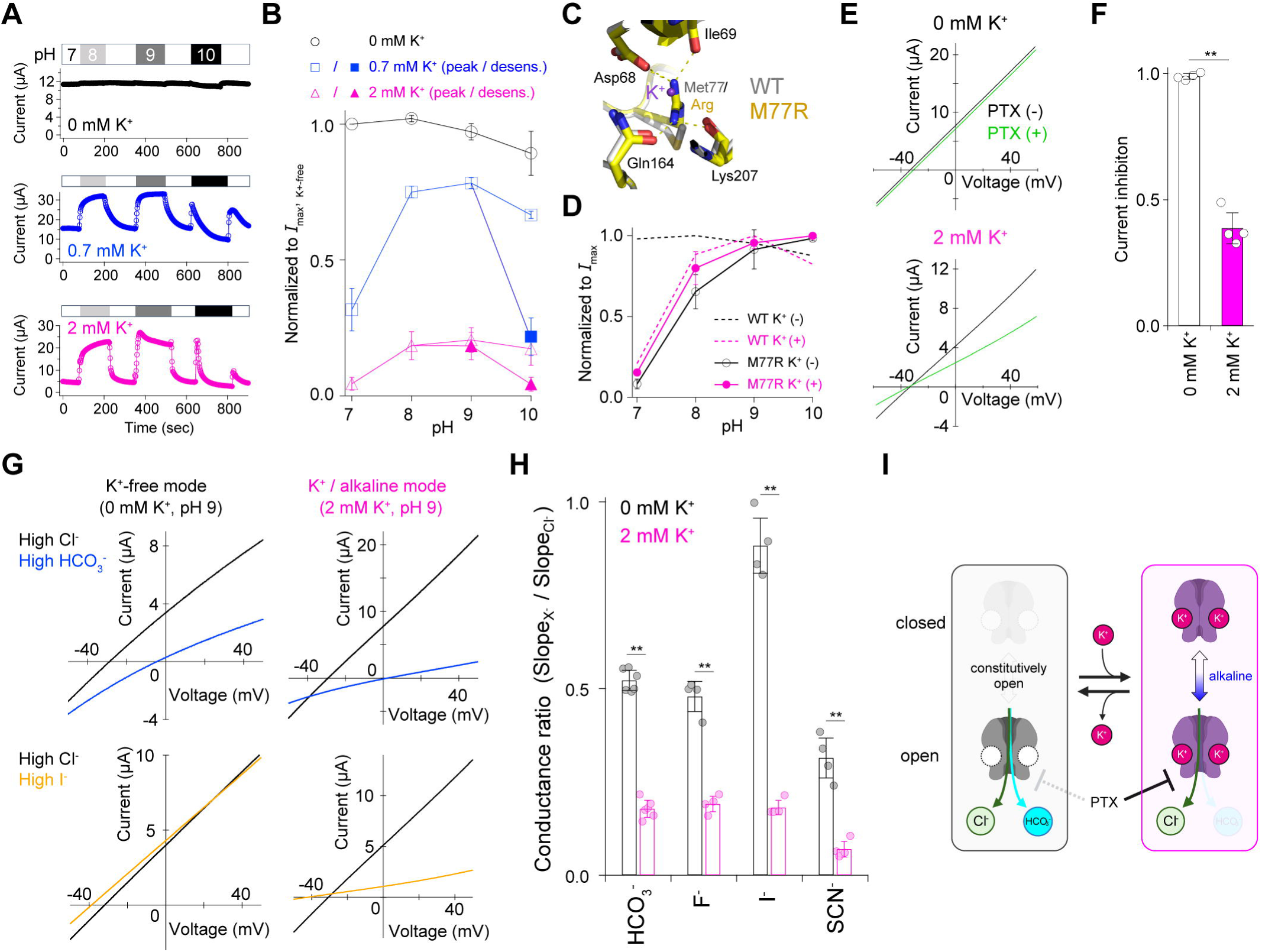
A K^+^-dependent mode-switching mechanism in DmAlka. (**A**) Representative time-courses of pH-dependence. Current amplitude at +60 mV, obtained from the ramp-pulse protocol, are used. (**B**) Plots of pH-dependence of the normalized current amplitude obtained from (A). The peak currents (open symbols) and steady-state / desensitized currents (filled symbols) are shown (n = 3-5). (**C**) Overlay of AF3-predicted structures of DmAlka WT (white) and M77R mutant (yellow), with the K^+^ binding residues shown as stick models. (**D**) Plots of pH-dependence of the normalized current amplitudes in M77R in the absence or presence of 2 mM KCl (n = 3). For comparison, the WT data in (B) normalized to their maximum amplitude are shown. (**E**) Representative current traces recorded in the absence (black) or presence (green) of 1 mM picrotoxin (PTX). (**F**) Bar plots comparing the PTX-induced current inhibition (n = 4-5). (G) Representative current traces obtained under the solutions with different major anions. (H) Plots of conductance ratios of various anions compared to Cl^□^, calculated from the slope values shown in (G) (n = 4-6). (**I**) A model of K^+^-dependent mode-switching in DmAlka. K^+^ binding switches the two modes with functional differences in both ligand (alkaline) sensitivity and pore properties.

We also found that the inhibitory effect of picrotoxin, a pore blocker of anion-channel Cys-loop receptors, on DmAlka was significantly reduced in WT in the absence of K^+^, and in the D82A mutant even in the presence of K^+^ (Fig. 3E, F, fig. S4F, G). These results indicate K^+^-induced conformational changes in the pore region. Therefore, the possibility that K^+^ modulates ion permeability and selectivity was investigated in detail. We evaluated the relative macroscopic conductance and selectivity for Cl^-^ versus other anions, of either K^+^ depletion-induced currents or alkaline-activated currents in the presence of K^+^. In the alkaline / K^+^ condition, Cl^-^ exhibited the highest conductance among the tested anions, as shown in the steepest slopes in the high Cl^-^ solutions, while this high Cl^-^ preference was significantly reduced in the K^+^-depleted condition (Fig. 3G, H, fig. S4H). In addition, the selectivity between tested anions determined by the shift of the reversal potentials was more pronounced in the presence of K^+^ than in the absence of K^+^ (Table S1). These results, together with those of picrotoxin, suggest that K^+^ binding induces a substantial conformational change in the pore region.

The K^+^ binding site is located at the ECD-TMD interface, which is critical for coupling ligand binding and gate opening in general Cys-loop receptors (Fig. 2A) (*18*). In particular, the K^+^ binding site is spatially connected, along the subunit interface, to the putative alkaline-sensing region that is positively charged for OH^-^ (fig. S2D). The K^+^ binding/unbinding could affect both these regions, through the rearrangements of the polar network at the subunit interface that surrounds the binding site. These observations indicate that K^+^ allosterically modulates conformations of a whole channel including both ECD and TMD, as if it functions as a “mode-switcher” of functionally and structurally distinct two modes (Fig. 3I).

### Human glycine receptor **α**2 has the potential to be K^+^ sensitive

A human homolog of DmAlka, GlyR α2 variant A (HsGlyRα2A), was previously shown to be activated by external Cs^+^ (*19*), suggesting its potential to be sensitive to extracellular K^+^. However, HsGlyRα2A WT did not show any K^+^ dependence (Fig. 4A, B). Therefore, we investigated the extent of the “trace” of K^+^-binding site in HsGlyRs by introducing mutations. We used a cyro-EM structure of HsGlyRα2/β heteromer (*20*) and aligned its two neighboring α2 subunits with those in the predicted structure of DmAlka. Based on the comparison, we designed a candidate mutant expected to have K^+^ sensitivity with quadruple mutations (Qm), composed of two mutations (S81D, P173Q) to coordinate K^+^ directly and two additional mutations (G83N, N95D) to strengthen the network of polar interactions observed in the DmAlka structure (Fig. 4C). Electrophysiological recordings of the HsGlyRα2A Qm showed a significantly larger basal currents in 98 mM high-K^+^ solution than the high-Na^+^ or NMDG^+^ solution, even without glycine application (Fig. 4A, B). These results indicate that HsGlyRα2A has the potential to be sensitive to K^+^ by just four mutations (Fig. 4D). It is noteworthy that, while K^+^ reduced the currents in DmAlka, in HsGlyRα2A Qm, K^+^ enhances basal currents and glycine-evoked activation occurs irrespective of the presence or the absence of K^+^ (Fig. 1A, 4A, 4D).

**Fig. 4.**
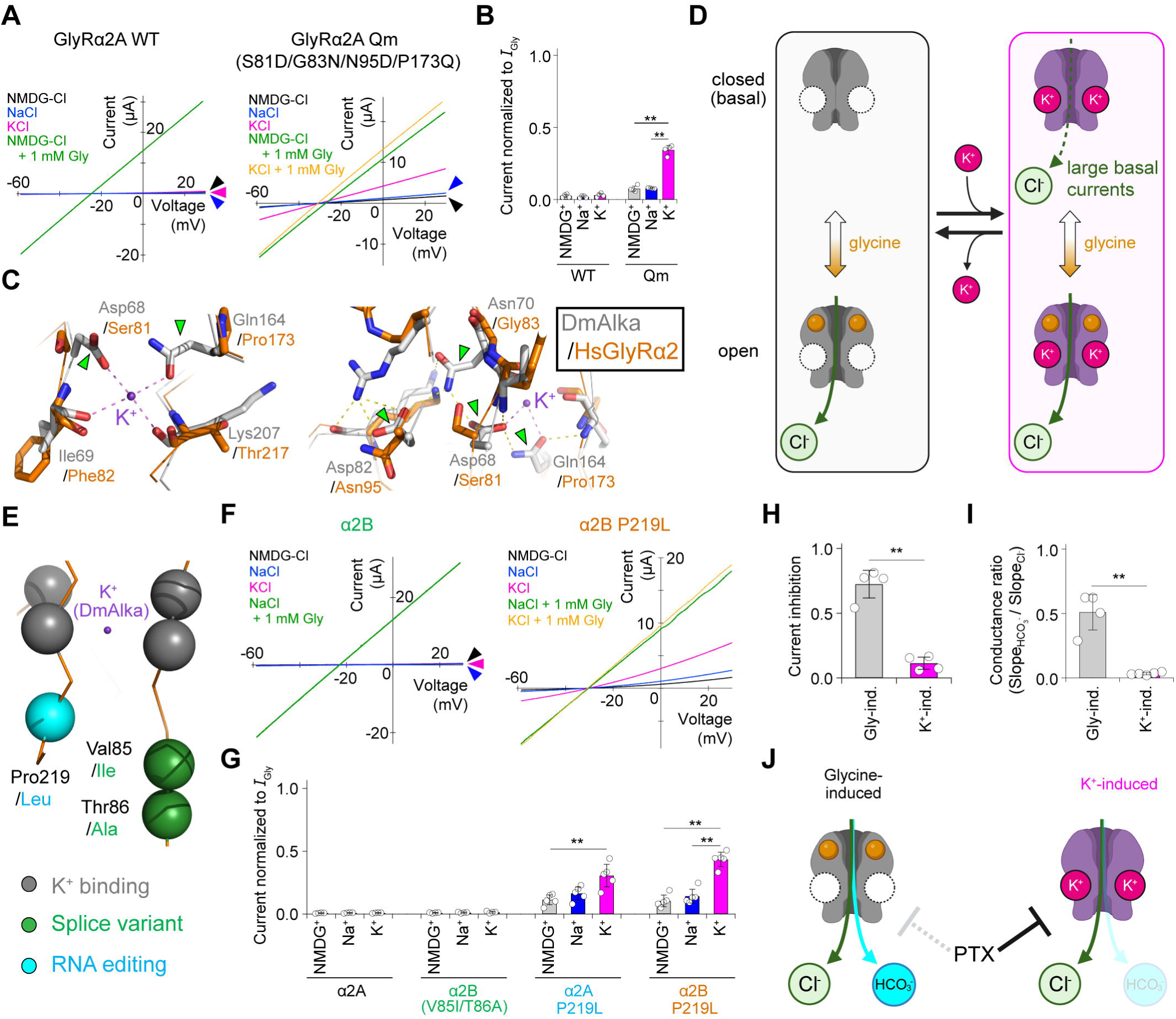
K^+^ sensitivity in HsGlyRα2 mutant and native variants. (**A**) Cation dependence of HsGlyRα2A WT (left) and Qm (right). (**B**) Plots of the normalized current amplitude under the different cations, shown as (A) (n = 4). (**C**) Structural comparison of DmAlka (white) and HsGlyRα2A (orange, PDB: 7L31). The K^+^- binding site (left) and its surrounding polar interactions (right) are highlighted. The yellow dashed lines indicate the polar interactions formed by the Qm mutations. Green arrowheads indicate the 4 Qm mutations. (**D**) A model of K^+^ dependence in HsGlyRα2A Qm and α2B P219L. K^+^ binding induces the larger basal currents in the absence of glycine. (**E**) Substitution positions of α2B and P219L in the structure of HsGlyR α2A (orange ribbon), superimposed with the K^+^ (purple) in that of DmAlka. Spheres indicate the C atoms of each position. (**F**) Cation dependence of HsGlyR α2B (left) and α2B P219L (right). (**G**) Plots of the normalized current amplitude under different cations, shown as (F). (**H**, **I**) Plots comparing PTX inhibition (n = 4) (H) and conductance ratio between high Cl^-^ and high HCO_3_^-^ (n = 4-5) (I) under glycine- and K^+^-induced modes. (**J**) A model illustrating the pore properties in glycine- and K^+^-induced modes.

We next investigated whether certain native GlyR variants might exhibit K^+^ sensitivity. In HsGlyRα2, two naturally occurring substitutions, the splice variant α2B (V85T/T86A) and the RNA-edited form (P219L), are known to enhance both glycine and Cs^+^ sensitivity (*19*, *21–23*). Notably, all these substitutions occur near the putative K^+^ binding site (Fig. 4E). We found that P219L native substitution conferred the K^+^- dependence to HsGlyRα2A, similar to the artificial Qm mutation (Fig. 4F, fig. S5A). The splice variant α2B itself did not exhibit a K^+^-induced current, but tended to enhance the K^+^- dependent increase in Cl^-^ current when combined with P219L (Fig. 4G). These results suggest that the structural deviation of the binding site caused by the substitutions surrounding it could confer K^+^ sensitivity. We also investigated whether K^+^ affects pore function and conformation in HsGlyRα2B P219L or not, similar to DmAlka. Inhibition by picrotoxin was more pronounced in K^+^-induced currents than in glycine-induced currents (Fig. 4H, fig. S5B). In addition, K^+^-induced currents exhibited significantly higher permeability and selectivity of Cl^-^ over HCO_3_ ^-^ than glycine-evoked currents (Fig. 4I, fig. S5C, Table S1). These effects of K^+^ on pore properties are almost identical to those observed in DmAlka (Fig. 3F, 3H). These results suggest, even though K^+^ exerts the opposite effects on the current amplitude, a decrease in DmAlka and an increase in HsGlyRα2B P219L respectively, both of them share the K^+^-induced mode-switching mechanism (Fig. 3I, 4J). In summary, native HsGlyRα2 retains the potential for K^+^- dependent mode-switching mechanism, analogous to DmAlka.

## Discussion

In this study, we characterized DmAlka as the first eukaryotic protein with a regulatory site for physiologically relevant concentrations of extracellular K^+^ and proposed a structural model for its selective K^+^ recognition. In addition, we found that the K^+^ binding in DmAlka induces the mode-switching mechanism, which is novel among Cys-loop receptor channels. These mechanisms are, at least partially, conserved even in the evolutionarily distant HsGlyRα2, indicating their broader and conserved roles across species.

Regarding the extracellular K^+^ sensitivity in life other than eukaryotes, bacteria have a unique kinase protein KdpD that detects extracellular K^+^ in µM order, and a spike protein of Hazara virus responds to tens of mM K^+^ on its cell infection (*24*, *25*). Many enzymes require K^+^ or Na^+^ with low affinities for their activities, as stable cofactors rather than as regulatory signals (*26*, *27*). In contrast, DmAlka has a clear extracellular binding site, and displayed IC_50_ of ∼1mM and strict K^+^ selectivity, with Na^+^ almost insensitive (Fig. 1B and fig. S1D). Its high K^+^ selectivity is plausibly explained by the predicted structure of K^+^ binding site (Fig. 2B). Discovery of the sensing module for extracellular K^+^ in DmAlka suggests the possibility of “extracellular K^+^ signaling” previously overlooked in eukaryotes. Similar to the case where DmAlka has been described merely as an alkaline-activated ion channel, other proteins may also harbor hidden K^+^-sensing machinery that has yet to be uncovered.

The effective range of K^+^ concentrations for DmAlka matches the physiological extracellular K^+^ concentrations in the *Drosophila* brain, which are in the millimolar range (Fig. 1A, B) (*28*). DmAlka has been reported to function in the brain (*9*), although less information is available regarding the role in brain than that of alkaline sensation (*10*, *29*). Interestingly, our search for public transcriptome databases and genetic approach to label the DmAlka expressing cells indicate that it is expressed in the brain glial cells, which play a major role in K^+^ clearance (fig. S6) (*30*). Given these observations, DmAlka may contribute to unidentified physiological mechanisms in the brain that rely on extracellular K^+^. For instance, the expression of DmAlka might give the glial cells the ability to modulate Cl^-^ conductance level in response to alterations in extracellular K^+^ concentration, thereby actively regulating K^+^ clearance.

In the broader superfamily of pentameric ligand-gated ion channels (pLGICs), including the Cys-loop receptor family, not only orthosteric ligands but also various allosteric modulators have been extensively investigated for each receptor (*31–35*). They include anesthetics, alcohols, synthetic compounds, as well as cations (*32*, *35*). Ca^2+^, Ba^2+^ and Cs^+^ are known to bind to the ECD-TMD interface in some bacterial pLGICs (*36–38*). In a bacterial extracellular Ca^2+^-gated channel, DeCLIC, its Ca^2+^ binding site is partly overlapped with the K^+^ binding site in DmAlka (fig. S7A, B) (*37*). Comparison of the structures of DeCLIC in apo and Ca^2+^-bound states shows a structural rearrangement of the loops in the ECD-TMD interface (fig. S7B), which could similarly occur for DmAlka on K^+^ binding to initiate its mode-switching mechanism. Ca^2+^ and Zn^2+^ are the allosteric modulators of nicotinic acetylcholine receptor α7 and HsGlyRs, respectively (fig. S7C-F) (*39*, *40*). Their binding sites are close to but distinct from those in DmAlka. These comparisons with other pLGIC structures show that the ECD-TMD interface in DmAlka forms a unique cation binding site among eukaryotic Cys-loop receptors, in which K^+^ binding regulates the pore opening through the structural rearrangements of the interfacial loops.

Furthermore, we newly found an unprecedented mechanism, in which the cation binding functions as a “mode-switcher” accompanying a global transition between two modes with differences in ligand-dependent activation, desensitization, blocker binding, and even ion permeability and selectivity (Fig. 3I and 4D, J). A large number of studies have shown that the conformational rearrangement in the ECD-TMD interface is critical for and coupled with the gate opening, as well as desensitization, in the TMDs (*18*, *41–44*). Since the desensitization is caused by structural rearrangements in the lower portion of the pore-lining M2 helices, the K^+^ binding at the ECD-TMD interface could affect the properties regulated by this region, which include ion selectivity and picrotoxin binding (43). In both DmAlka and HsGlyRα2B P219L, other anions than Cl^-^ permeates in larger conductance with lower selectivity, as well as more reduced picrotoxin binding, in K^+^- unbound states than K^+^-bound states (Figs. 3F, 3H, 4H, 4I), as if the pore becomes “wider and looser”. Interestingly, the changes in these functional properties seem to resemble the structural transition between the normal and expanded open states observed in cryo-EM structures of glycine receptors in a lipidic environment (41), in which the lower part of M2 helices was wider in the expanded state than in the normal state. The similar mechanism might occur on K^+^-dependent mode-switching in DmAlka. It is notable from a physiological point of view, related to the DmAlka expression in glial cells, that these “wider and looser” pore allows the permeation of HCO_3_ ^-^ much more in the K^+^-unbound state than the K^+^-bound state (Fig. 3H, 4I). In glial cells, HCO_3_ ^-^ efflux is known to buffer extracellular pH during neuronal activity (*45*). The channel may allow HCO_3_ ^-^ efflux to facilitate pH buffering in the K^+^-unbound mode, while in the K^+^-bound mode, it may induce the more polarized potential reflecting the Cl^-^ gradient across the membranes.

It is noteworthy that, despite the considerable evolutionary distance from DmAlka, the variant of its human homolog still retains K^+^ sensitivity (Fig. 4G). The effective K^+^ concentration for HsGlyRα2B P219L is too high to show its potentiation (fig. S5D, E), considering the typical extracellular K^+^ concentration in the mammalian brains. However, the investigations of the extracellular K^+^ concentration in rats showed that it can rise over 50 or 80 mM during ischemic conditions (*46*, *47*). Interestingly, in the hippocampus isolated from human patients with temporal lobe epilepsy (TLE) who suffer from chronic neuronal firing, the relative expression of the α2B splice variant and the P219L RNA-edited form increased significantly compared to the α2A variant (*48*, *49*). This raises the possibility that these variants may contribute to either the TLE suppression or progression through the increased Cl^-^ conductance caused by high concentrations of K^+^. K^+^ sensitivity in the HsGlyR may serve as functional mechanisms under pathologically K^+^-accumulated conditions. Rb^+^ exhibited a similar effect to K^+^ even in lower concentrations (fig. S5D, E). Therefore, it could serve as a lead for the development of small-molecule modulators targeting the ECD-TMD interface. Such compounds would be valuable to investigate the possible physiological contribution of the mode-switching mechanism, and to develop therapeutics targeting glycine receptors with a fundamentally distinct principle from canonical drugs.

## Acknowledgement

We thank C. Naito and T. Yamamoto for technical assistance and all members of the Kubo laboratory for helpful discussions.

## Author contributions

M.S. and Y.K. conceived and supervised the project. T.S. and Y.S. designed and performed the experiments. T.S. performed and analyzed the electrophysiological data. Y.S. performed phylogenetic analysis and immunohistochemistry. T.S. and Y.S. prepared the figures. T.S. wrote the draft of the manuscript with the support from Y.S. T.S., Y.K., M.S., and Y.S. completed the manuscript. All authors approved the final version of the manuscript.

## Data and materials availability

All the data supporting this study are available within this article and supplementary data or the online repository.

## References

1. J. Stautz, Y. Hellmich, M. F. Fuss, J. M. Silberberg, J. R. Devlin, R. B. Stockbridge, I. Hänelt, Molecular mechanisms for bacterial potassium homeostasis. J. Mol. Biol. 433, 166968 (2021).

2. A. A. McDonough, R. A. Fenton, Potassium homeostasis: sensors, mediators, and targets. Pflugers Arch. 474, 853–867 (2022).

3. A. Danchin, P. I. Nikel, Why nature chose potassium. J. Mol. Evol. 87, 271–288 (2019).

4. A. A. McDonough, J. H. Youn, Potassium homeostasis: The knowns, the unknowns, and the health benefits. Physiology (Bethesda) 32, 100–111 (2017).

5. U. K. Udensi, P. B. Tchounwou, Potassium homeostasis, Oxidative stress, and human disease. Int. J. Clin. Exp. Physiol. 4, 111–122 (2017).

6. D. Podar, F. J. M. Maathuis, Primary nutrient sensors in plants. iScience 25, 104029 (2022).

7. R.-J. Tang, F.-G. Zhao, Y. Yang, C. Wang, K. Li, T. J. Kleist, P. G. Lemaux, S. Luan, A calcium signalling network activates vacuolar K+ remobilization to enable plant adaptation to low-K environments. Nat. Plants 6, 384–393 (2020).

8. M. W. Szczerba, D. T. Britto, H. J. Kronzucker, K+ transport in plants: physiology and molecular biology. J. Plant Physiol. 166, 447–466 (2009).

9. L. Frenkel, N. I. Muraro, A. N. Beltrán González, M. S. Marcora, G. Bernabó, C. Hermann-Luibl, J. I. Romero, C. Helfrich-Förster, E. M. Castaño, C. Marino-Busjle, D. J. Calvo, M. F. Ceriani, Organization of circadian behavior relies on glycinergic transmission. Cell Rep. 19, 72–85 (2017).

10. T. Mi, J. O. Mack, W. Koolmees, Q. Lyon, L. Yochimowitz, Z.-Q. Teng, P. Jiang, C. Montell, Y. V. Zhang, Alkaline taste sensation through the alkaliphile chloride channel in Drosophila. Nat Metab 5, 466–480 (2023).

11. L. A. Gay, P. R. Stanfield, The selectivity of the delayed potassium conductance of frog skeletal muscle fibers. Pflugers Arch. 378, 177–179 (1978).

12. J. Abramson, J. Adler, J. Dunger, R. Evans, T. Green, A. Pritzel, O. Ronneberger, L. Willmore, A. J. Ballard, J. Bambrick, S. W. Bodenstein, D. A. Evans, C.-C. Hung, M. O’Neill, D. Reiman, K. Tunyasuvunakool, Z. Wu, A. Žemgulytė, E. Arvaniti, C. Beattie, O. Bertolli, A. Bridgland, A. Cherepanov, M. Congreve, A. I. Cowen-Rivers, A. Cowie, M. Figurnov, F. B. Fuchs, H. Gladman, R. Jain, Y. A. Khan, C. M. R. Low, K. Perlin, A. Potapenko, P. Savy, S. Singh, A. Stecula, A. Thillaisundaram, C. Tong, S. Yakneen, E. D. Zhong, M. Zielinski, A. Žídek, V. Bapst, P. Kohli, M. Jaderberg, D. Hassabis, J. M. Jumper, Accurate structure prediction of biomolecular interactions with AlphaFold 3. Nature 630, 493–500 (2024).

13. A. J. Thompson, H. A. Lester, S. C. R. Lummis, The structural basis of function in Cys-loop receptors. Q. Rev. Biophys. 43, 449–499 (2010).

14. P.-J. Corringer, F. Poitevin, M. S. Prevost, L. Sauguet, M. Delarue, J.-P. Changeux, Structure and pharmacology of pentameric receptor channels: from bacteria to brain. Structure 20, 941–956 (2012).

15. H. A. Lester, M. I. Dibas, D. S. Dahan, J. F. Leite, D. A. Dougherty, Cys-loop receptors: new twists and turns. Trends Neurosci. 27, 329–336 (2004).

16. J. Mähler, I. Persson, A study of the hydration of the alkali metal ions in aqueous solution. Inorg. Chem. 51, 425–438 (2012).

17. Y. Zhou, J. H. H. Morais-Cabral, A. Kaufman, R. MacKinnon, Chemistry of ion coordination and hydration revealed by a K channel-Fab complex at 2.0 Å resolution. Nature 414, 43–48 (2001).

18. M. Bartos, J. Corradi, C. Bouzat, Structural basis of activation of cys-loop receptors: the extracellular-transmembrane interface as a coupling region. Mol. Neurobiol. 40, 236–252 (2009).

19. S. Fricke, M. Harnau, F. Hetsch, H. Liu, J. Leonhard, A. Eylmann, P. Knauff, H. Sun, M. Semtner, J. C. Meier, Cesium activates the neurotransmitter receptor for glycine. Front. Mol. Neurosci. 16 (2023).

20. H. Yu, X.-C. Bai, W. Wang, Characterization of the subunit composition and structure of adult human glycine receptors. Neuron 109, 2707–2716.e6 (2021).

21. J. C. Meier, C. Henneberger, I. Melnick, C. Racca, R. J. Harvey, U. Heinemann, V. Schmieden, R. Grantyn, RNA editing produces glycine receptor alpha3(P185L), resulting in high agonist potency. Nat. Neurosci. 8, 736–744 (2005).

22. S. A. Eichler, B. Förstera, B. Smolinsky, R. Jüttner, T.-N. Lehmann, M. Fähling, G. Schwarz, P. Legendre, J. C. Meier, Splice-specific roles of glycine receptor alpha3 in the hippocampus. Eur. J. Neurosci. 30, 1077–1091 (2009).

23. P. S. Miller, R. J. Harvey, T. G. Smart, Differential agonist sensitivity of glycine receptor alpha2 subunit splice variants: Glycine receptorα2 subunit spice variants. Br. J. Pharmacol. 143, 19–26 (2004).

24. E. K. Punch, S. Hover, H. T. W. Blest, J. Fuller, R. Hewson, J. Fontana, J. Mankouri, J. N. Barr, Potassium is a trigger for conformational change in the fusion spike of an enveloped RNA virus. J. Biol. Chem. 293, 9937–9944 (2018).

25. V. Laermann, E. Ćudić, K. Kipschull, P. Zimmann, K. Altendorf, The sensor kinase KdpD of Escherichia coli senses external K+: KdpD senses external K+. Mol. Microbiol. 88, 1194–1204 (2013).

26. M. J. Page, E. Di Cera, Role of Na+ and K+ in enzyme function. Physiol. Rev. 86, 1049–1092 (2006).

27. E. Di Cera, A structural perspective on enzymes activated by monovalent cations. J. Biol. Chem. 281, 1305–1308 (2006).

28. G. A. B. Armstrong, E. C. Rodríguez, R. Meldrum Robertson, Cold hardening modulates K+ homeostasis in the brain of Drosophila melanogaster during chill coma. J. Insect Physiol. 58, 1511–1516 (2012).

29. P. Pandey, B. Shrestha, Y. Lee, Avoiding alkaline taste through ionotropic receptors. iScience 27, 110087 (2024).

30. A. Bellot-Saez, O. Kékesi, J. W. Morley, Y. Buskila, Astrocytic modulation of neuronal excitability through K+ spatial buffering. Neurosci. Biobehav. Rev. 77, 87–97 (2017).

31. Y. Chang, Y. Huang, P. Whiteaker, Mechanism of allosteric modulation of the cys-loop receptors. Pharmaceuticals (Basel) 3, 2592–2609 (2010).

32. C. F. Burgos, G. E. Yévenes, L. G. Aguayo, Structure and pharmacologic modulation of inhibitory Glycine receptors. Mol. Pharmacol. 90, 318–325 (2016).

33. U. Breitinger, H.-G. Breitinger, Modulators of the inhibitory Glycine receptor. ACS Chem. Neurosci. 11, 1706–1725 (2020).

34. A. Taly, J. Hénin, J.-P. Changeux, M. Cecchini, Allosteric regulation of pentameric ligand-gated ion channels: an emerging mechanistic perspective. Channels (Austin) 8, 350–360 (2014).

35. L. Sauguet, A. Shahsavar, M. Delarue, Crystallographic studies of pharmacological sites in pentameric ligand-gated ion channels. Biochim. Biophys. Acta 1850, 511–523 (2015).

36. I. Zimmermann, A. Marabelli, C. Bertozzi, L. G. Sivilotti, R. Dutzler, Inhibition of the prokaryotic pentameric ligand-gated ion channel ELIC by divalent cations. PLoS Biol. 10, e1001429 (2012).

37. H. Hu, R. J. Howard, U. Bastolla, E. Lindahl, M. Delarue, Structural basis for allosteric transitions of a multidomain pentameric ligand-gated ion channel. Proc. Natl. Acad. Sci. U. S. A. 117, 13437–13446 (2020).

38. H. Hu, Á. Nemecz, C. Van Renterghem, Z. Fourati, L. Sauguet, P.-J. Corringer, M. Delarue, Crystal structures of a pentameric ion channel gated by alkaline pH show a widely open pore and identify a cavity for modulation. Proc. Natl. Acad. Sci. U. S. A. 115, E3959–E3968 (2018).

39. K. Kindig, E. Gibbs, D. Seiferth, P. C. Biggin, S. Chakrapani, Mechanisms underlying modulation of human GlyRα3 by Zn2+ and pH. Sci. Adv. 10, eadr5920 (2024).

40. C. M. Noviello, A. Gharpure, N. Mukhtasimova, R. Cabuco, L. Baxter, D. Borek, S. M. Sine, R. E. Hibbs, Structure and gating mechanism of the α7 nicotinic acetylcholine receptor. Cell 184, 2121–2134.e13 (2021).

41. J. Yu, H. Zhu, R. Lape, T. Greiner, J. Du, W. Lü, L. Sivilotti, E. Gouaux, Mechanism of gating and partial agonist action in the glycine receptor. Cell 184, 957–968.e21 (2021).

42. J. Du, W. Lü, S. Wu, Y. Cheng, E. Gouaux, Glycine receptor mechanism elucidated by electron cryo-microscopy. Nature 526, 224–229 (2015).

43. A. Kumar, S. Basak, S. Rao, Y. Gicheru, M. L. Mayer, M. S. P. Sansom, S. Chakrapani, Mechanisms of activation and desensitization of full-length glycine receptor in lipid nanodiscs. Nat. Commun. 11, 3752 (2020).

44. S. Shi, S. N. Lefebvre, L. Peverini, A. H. Cerdan, P. Milán Rodríguez, M. Gielen, J.-P. Changeux, M. Cecchini, P.-J. Corringer, Illumination of a progressive allosteric mechanism mediating the glycine receptor activation. Nat. Commun. 14, 795 (2023).

45. S. M. Theparambil, P. S. Hosford, I. Ruminot, O. Kopach, J. R. Reynolds, P. Y. Sandoval, D. A. Rusakov, L. F. Barros, A. V. Gourine, Astrocytes regulate brain extracellular pH via a neuronal activity-dependent bicarbonate shuttle. Nat. Commun. 11, 5073 (2020).

46. A. J. Hansen, The extracellular potassium concentration in brain cortex following ischemia in hypo- and hyperglycemic rats. Acta Physiol. Scand. 102, 324–329 (1978).

47. M. Müller, G. G. Somjen, Na(+) and K(+) concentrations, extra- and intracellular voltages, and the effect of TTX in hypoxic rat hippocampal slices. J. Neurophysiol. 83, 735–745 (2000).

48. S. A. Eichler, S. Kirischuk, R. Jüttner, P. K. Schaefermeier, P. Legendre, T.-N. Lehmann, T. Gloveli, R. Grantyn, J. C. Meier, Glycinergic tonic inhibition of hippocampal neurons with depolarizing GABAergic transmission elicits histopathological signs of temporal lobe epilepsy. J. Cell. Mol. Med. 12, 2848–2866 (2008).

49. S. Kankowski, B. Förstera, A. Winkelmann, P. Knauff, E. E. Wanker, X. A. You, M. Semtner, F. Hetsch, J. C. Meier, A novel RNA editing sensor tool and a specific agonist determine neuronal protein expression of RNA-edited Glycine receptors and identify a genomic APOBEC1 dimorphism as a new genetic risk factor of epilepsy. Front. Mol. Neurosci. 10, 439 (2017).

