## Supplementary Files for "Extracellular K^+^ modulates the pore conformations of Cys-loop receptor anion channels"

**The PDF file includes:**

Materials and Methods

Figs. S1 to S7

Tables S1

References

**Materials and Methods**

Ethical Approval

All animal experiments using *Xenopus* oocytes were approved by and conducted in accordance with the guidelines of the Animal Care Committee of the National Institutes of Natural Sciences (umbrella institution of the National Institute for Physiological Sciences, Japan).

Molecular biology, cRNA Preparation and Injection into *Xenopus laevis* Oocytes

The cDNA of *Drosophila melanogaster* CG12344 (DmAlka, NCBI Reference Sequence: NP_001286297.1) was synthesized *de novo* (eurofins) and incorporated into the pGEMHE plasmid using the EcoRI restriction site. The cDNA of human glycine receptor α2A (HsGlyRα2A, NP_001112357) was purchased (GenScript) and incorporated into the pGEMHE plasmid using SmaI site. Preparation and injection of cRNAs into *Xenopus* oocytes were performed as previously described in detail (*1*). In brief, the cRNAs were transcribed using mMESSAGE mMACHINE T7 kit (Thermo Fisher Scientific) from the linearized cDNAs and stored at -80°C until use. The oocytes were collected from *Xenopus laevis* (Xenopus Yoshoku Kyouzai) by surgical operation under anesthesia with 0.15% tricaine. Defolliculated *Xenopus* oocytes were obtained by enzymatic treatment using collagenase (Sigma-Aldrich), and maintained at 17°C in frog Ringer's solution (in mM: 88 NaCl, 1 KCl, 2.4 $NaHCO_3$, 0.3 Ca $(NO_3)_2$, 0.41 $CaCl_2$ and 0.82 $MgSO_4$, 15 HEPES-NaOH (pH7.6)) with 0.1% penicillin–streptomycin (Sigma-Aldrich). Oocytes were injected with 50 nl of cRNA solution using a Nanoject II microinjector (Drummond Scientific Company). The concentrations of DmAlka cRNAs were adjusted for injection depending on the experimental condition: 0.001–0.05 µg/µL for $K^+$-free conditions, and 0.1-0.5 µg/µL for conditions in the presence of $K^+$. HsGlyRα2A cRNAs were injected at concentrations ranging from 0.01-0.1 µg/µL. Currents were measured 1 to 4 days after injection, depending on the current amplitude.

<u>Electrophysiological Recordings</u>

Two-electrode voltage-clamp recordings using *Xenopus* oocytes were performed at room temperature using an OC-725C amplifier (Warner Instruments) and pClamp11 software (Molecular Devices). Data were digitized at 10 kHz through Digidata1440 (Molecular Devices). The resistance of Borosilicate glass microelectrodes was adjusted to be 0.2–0.5MΩ when filled with the solution of 3M K-acetate and 10mM KCl.

For the experiments in DmAlka, unless especially mentioned here or in each figure legend, the standard solution was ND96 (in mM: 96 NaCl, 2 KCl, 1.8 $CaCl_2$, 1 $MgCl_2$ and 5 HEPES-NaOH (pH 7.4)). To explore the effects of $K^+$ on DmAlka currents, 2 mM KCl in the solutions was generally replaced with 2 mM NMDG-Cl. Adjustment of pH to investigate alkaline-dependence was achieved by adding 5 mM HEPES-NaOH for pH 7.0 and 8.0, and 5 mM CAPSO-NaOH for pH 9.0 and 10.0, instead of HEPES-NaOH (pH 7.4). To investigate the effects of PTX or anion exchanges under $K^+$/alkaline-mode, the oocytes were incubated in the solution with pH 7.0 for more than 30 min., then voltage-clamped, and perfused with the solution with pH 9.0 for long enough to reach at peak amplitude, followed by the rapid solution exchanges with those containing 1 mM PTX or different anions. The high $F^-$, $I^-$,

$SCN^-$ solutions contain each 96 mM sodium anion instead of 96 mM NaCl in the standard solution. The $K^+$ free-mode was investigated with the solutions containing 2 mM NMDG-Cl, instead of KCl, at pH9.0, to maintain the experimental consistency.

For the experiments in HsGlyRα2 series, unless especially mentioned here or in each figure legend, the standard solution was as follows; in mM: 98 NaCl (or 98 NMDG-Cl), 1.8 $CaCl_2$, 1 $MgCl_2$ and 5 HEPES-NMDG (pH 7.4). To investigate the effects of cations on HsGlyRα2 series, 98 mM NaCl or NMDG-Cl in the standard solutions was replaced with 98 mM KCl or 98 mM RbCl. The evoked current amplitudes were normalized to those by 1 mM glycine. Dose-dependence of $K^+$ and $Rb^+$ was examined using the solutions containing a total of 98 mM XCl, where $X^+$ represents the sum of $Na^+$ and either $K^+$ or $Rb^+$. To investigate the effects of picrotoxin (PTX) or anion exchanges under $K^+$-induced mode, the oocytes were voltage-clamped in the standard solution, and fully replaced with the solution containing 98 mM KCl, followed by the rapid solution exchanges with those containing 10 μM PTX or different anions. The high $HCO_3^-$ solution contains 98 mM $NaHCO_3$ or $KHCO_3$, instead of 98 mM NaCl or NMDG-Cl, in the standard solutions. The glycine-induced mode was evoked by 0.1 mM glycine in the standard solution, followed by the solution exchanges as rapid as possible to minimize the effects of slow, but steadily accumulating desensitization.

For the experiments using the solutions containing high concentration of anions, the salt bridge, which are composed of 3% agar and either 1M NaCl in DmAlka or 1M KCl in HsGlyRα2A, were used to minimize the shift in liquid junction potentials caused by lower mobility of certain anions. In addition, to further minimize the residual shift that occur despite of its usage (at most ~5 mV), the voltage offset caused by solution exchanges was measured in advance with the electrodes soaked in the solutions before the oocyte recordings, and used to correct the I-V relationships in the subsequent data analysis. All representative traces shown as figures are corrected versions. For the experiments using $HCO_3^-$, freshly prepared $NaHCO_3$ or $KHCO_3$ solution was made immediately before the experiments to avoid their degradation. In HsGlyR α2B P219L, for the anion exchange experiments using $HCO_3^-$, the pH of the solutions containing either 98 mM NaCl or KCl was adjusted to pH 8.0-8.2 with NaOH, to align with the pH resulting from the simple inclusion of 96 mM $NaHCO_3$ or $KHCO_3$ in the standard solution.

Data Analysis and Statistics

Dose-response curves of the $K^+$-dependent inhibition were fitted with the Hill equation as follows:

$$I = I_{max} - \frac{(I_{max} - I_{min})}{1 + \left(\frac{IC_{50}}{[K^+]_{ext.}}\right)^n}$$

where $I$, $I_{max}$ and $I_{min}$ indicate the current amplitude at a certain concentration, maximal amplitude, and minimum amplitude respectively, n and $IC_{50}$ refer to the Hill slope and the half maximal inhibitory concentration respectively.

Ion selectivity was estimated by measuring the shifts in reversal potential induced by exchange between two bath solutions containing different anion species, under the assumption that only anions permeate to DmAlka and HsGlyRα2. Relative permeability ratios were calculated using the Goldman-Hodgkin-Katz equation as follows:

$$\Delta E_{rev} = E_{rev,B} - E_{rev,A} = \frac{RT}{zF} \ln \frac{P_{Cl^-_B} [Cl^-]_B + P_{X^-_B} [X^-]_B}{P_{Cl^-_A} [Cl^-]_A + P_{X^-_A} [X^-]_A}$$

where A and B indicate the bath solutions exchanged, $E_{rev, A}$ and $E_{rev, B}$ indicate the reversal potential measured under the solutions A and B, and $\Delta E_{rev}$ indicate the difference between $E_{rev, A}$ and $E_{rev, B}$, $P_{Cl-}$ and $P_{X-}$ indicate the permeability of $Cl^-$ and $X^-$ (each tested anion), $[Cl^-]$ and $[X^-]$ indicate the concentrations of $Cl^-$ and $X^-$ contained in each bath solution, R, T, z, and F refer to the gas constant, absolute temperature, charge of anions, and Faraday's constant, respectively.

For comparison of relative macroscopic conductance, I-V relationships obtained by the ramp-pulse protocols were fitted by linear function. The slope values obtained in each trace indicates a macroscopic anion conductance, and the ratio of the two slope values obtained under the two solutions containing different anions represents change in conductance caused by differences in anion species.

Data were analyzed using Igor Pro (WaveMetrics) and Microsoft Excel. All data are shown as mean ± S.D. For statistical comparisons, unpaired t-tests were used for two groups, One-way ANOVA followed by Dunnett's post hoc test for multiple comparisons against a control, and One-way ANOVA followed by Tukey's HSD test for intergroup comparisons. Significance levels were indicated as follows: $*p < 0.05$, $**p < 0.01$, and $p \geqq 0.05$ for not significant (n.s.).

AlphaFold3 Structure Prediction and Structure Visualization

Structural prediction was performed using the AlphaFold server (https://alphafoldserver.com/) powered by AlphaFold 3 (*2*). Amino acid sequence of the full-length DmAlka and its homologs, including their signal peptide parts, was used to generate its homo-pentameric models. To estimate the $K^+$ or $Na^+$ binding sites, structures were generated by a query that included five cations in addition to protein sequences. The predicted structures for apo, $Na^+$-bound, and $K^+$-bound states in DmAlka showed only minimal differences (fig. S2E). Among the five predicted structures in each prediction trial, the one with the highest confidence score was used. The average distance between $K^+$ and coordinated oxygens was calculated from the values in all the five $K^+$, each of which showed a slight variation. All residue numbering in HsGlyRα2 and DmAlka corresponds to the full-length sequence, including their signal peptides, rather than omitting it, in order to maintain consistency with the sequence numbering in the previous report of DmAlka (*10*). Visualizations of the structures were performed using PyMOL (Schrödinger, LLC). Electrostatic surface potential maps were obtained from the AF3-predicted output file using the PDB2PQR server (https://server.poissonboltzmann.org) (*3*) and visualized by PyMOL APBS Tool2.1 plugin.

Fly strains

Flies were raised on standard medium at 25°C on a 12:12 L/D cycle. The following fly stocks were used for immunohistochemistry: *CG12344(alka)-T2A-Gal4* (gifted by Dr. Shu Kondo), *10xUAS-IVS-mCD8::GFP* (BDSC #32185).

Immunohistochemistry

$CO_2$ anesthetized females were dissected in PBS. Dissected brains were fixed with 4% paraformaldehyde for 15 min at room temperature, blocked for 1h in PBS with 0.3% Triton X-100 and 10% ImmunoBlock (KAC Co., Ltd.), incubated for 1 day each (with washing in between) at room temperature in blocking solutions with a primary or secondary antibody. The brain was mounted in SeeDB2 (FUJIFILM Wako) to increase tissue transparency. We used the following primary and secondary antibodies at the indicated dilutions: 1:1000 chicken anti-GFP (abcam) and, 1:500 Alexa Fluor Plus 488 goat anti-chicken lgY (Life Technologies). Confocal images were acquired with Zeiss LMS900 confocal laser scanning microscope under 40x magnification.

Phylogenetic analysis of DmAlka

Putative homologs of DmAlka were searched by the protein local alignment search tool (BLASTP) analysis using the amino acid sequence of CG12344/DmAlka as the query for the National Center for Biotechnology information (NCBI) database (https://blast.ncbi.nlm.nih.gov/Blast.cgi). When multiple candidates of the homologous proteins were obtained from the same animal species, the one with the highest sequence identity was selected.

The phylogenetic tree was inferred using the Minimum Evolution (ME) method (*4*). The evolutionary distances were computed using the Poisson correction method and are in the units of the number of amino acid substitutions per site. The ME tree was searched using the Close-Neighbor-Interchange (CNI) algorithm. The Neighbor-joining algorithm was used to generate the initial tree. The pairwise deletion option was applied to all ambiguous positions for each sequence pair resulting in a final data set comprising 750 positions. Phylogenetic analyses were conducted in MEGA12 software (*5*). All the sequences and predicted structures of the homologs are deposited in Dryad repository.

Data availability

All the data supporting this study are available within this article and supplementary data or the Dryad repository.

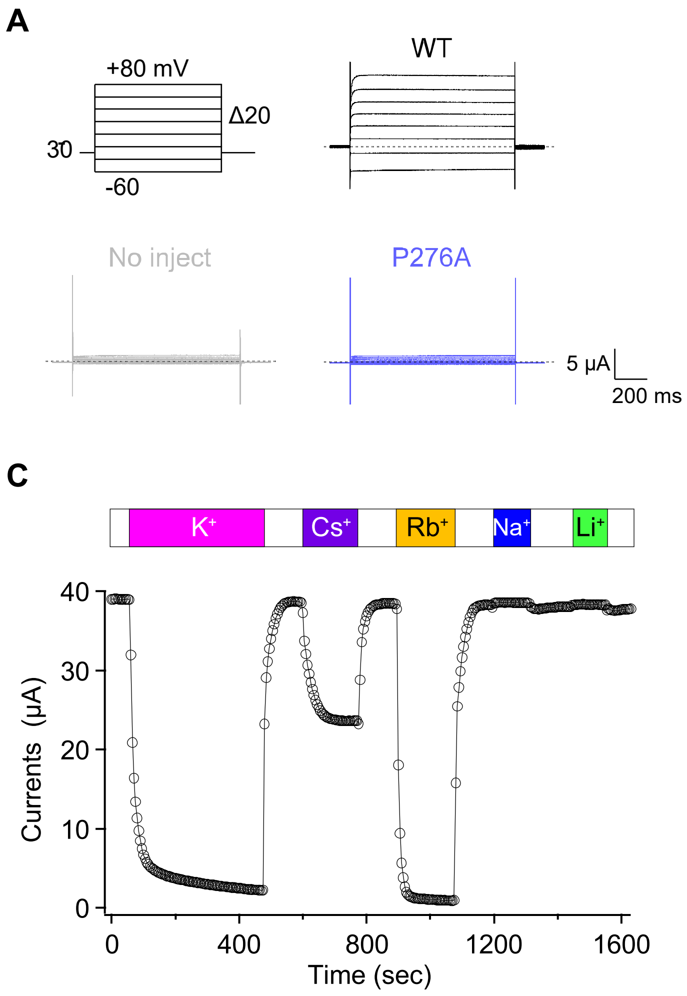


**Fig. S1. Characterization and cation dependence of DmAlka $Cl^-$ currents in *Xenopus* oocytes.**

185 (**A**) Representative current traces recorded from the oocytes injected with cRNAs of WT (black), P276A, a pore-dead mutant used in the previous report (*10*) (blue), or without cRNA (non-injected, grey) in the standard ND96 solution. The currents were elicited by the step pulse shown in the top left. (**B**) Current–voltage (I-V) relationships obtained from the recordings shown in (A) (n = 5-10). (**C**) A representative plot showing the time-dependent

change in the DmAlka current on exchanging the bath solutions from the basic 96 mM NMDG-Cl solutions to those containing 2 mM of each monovalent cation chloride. The current amplitude at +60 mV obtained by the ramp-pulse protocol from -60 mV to +60 mV every 5 sec was used and plotted. (**D**) Representative current traces under the respective bath solutions containing the same concentrations of $Cl^-$ and different concentrations of $Na^+$, $K^+$, or $NMDG^+$ as that of ND96.

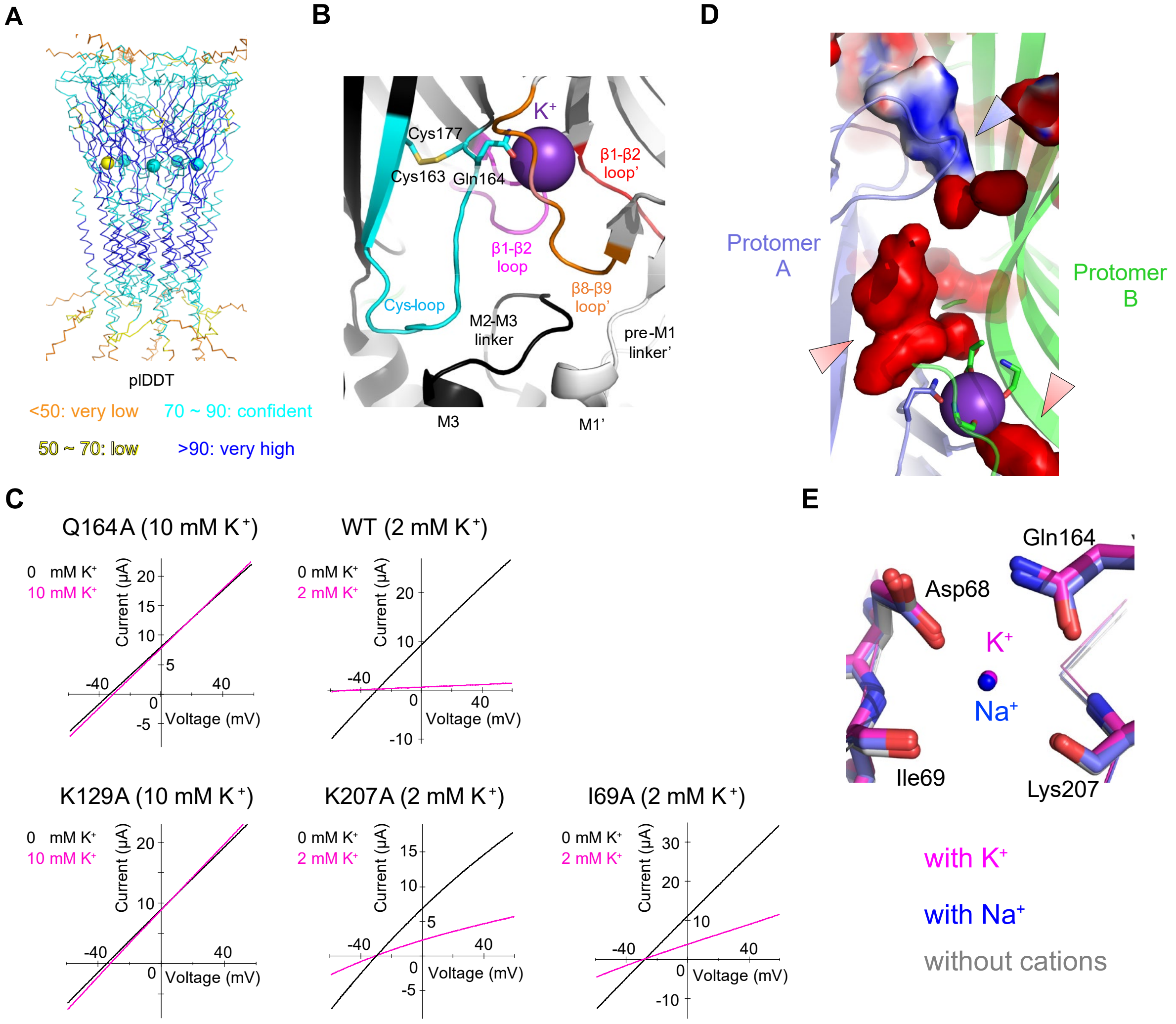


**Fig. S2. Structural features of the K$^+$ binding site in an AlphaFold 3-predicted model of DmAlka.**

(**A**) A whole view of the structural model of a full-length DmAlka pentamer bound with K$^+$ predicted by AlphaFold 3 (AF3). The model is colored according to the pLDDT scores as shown at the bottom. Spheres indicate the bound K$^+$. (**B**) A close-up view of the K$^+$ binding site and its surrounding regions. K$^+$ is depicted as a purple sphere. The side chains of a disulfide bond between two cysteines characteristic in Cys-loop receptors and its neighboring Gln164, which interacts directly with $K^+$, are shown as stick models. Two different protomers are colored in black and white, respectively. The Cys-loop, β1–β2 loops, and β8–β9 loop in two neighboring protomers are colored differently. The regions labeled with an apostrophe indicates that they belong to a different protomer from that without it. (**C**) Representative current traces recorded from the oocytes expressing DmAlka WT or mutants (I69A K129A, Q164A, K207A). Currents were recorded under the solutions without (black), or with 2 or 10 mM (magenta) concentrations of $K^+$. (**D**) The electrostatic surface potential of the region between the neighboring two protomers (pale blue and green), focusing on the $K^+$ binding site and a putative alkaline-sensing region. The surface potentials are contoured and colored from -5 kT (red) to +5 kT (blue). Positively charged regions around the $K^+$ binding sites and negatively charged alkaline-sensitive site are indicated by red and blue arrowheads, respectively. (**E**) Structural alignment of three AF3-predicted structures of the $K^+$ binding site in the $K^+$-bound state (magenta), $Na^+$-bound state (blue), and apo state (gray). The side chains of four key residues directly involved in $K^+$ recognition (Asp68, Ile69, Gln164, and Lys207) are shown as stick models.

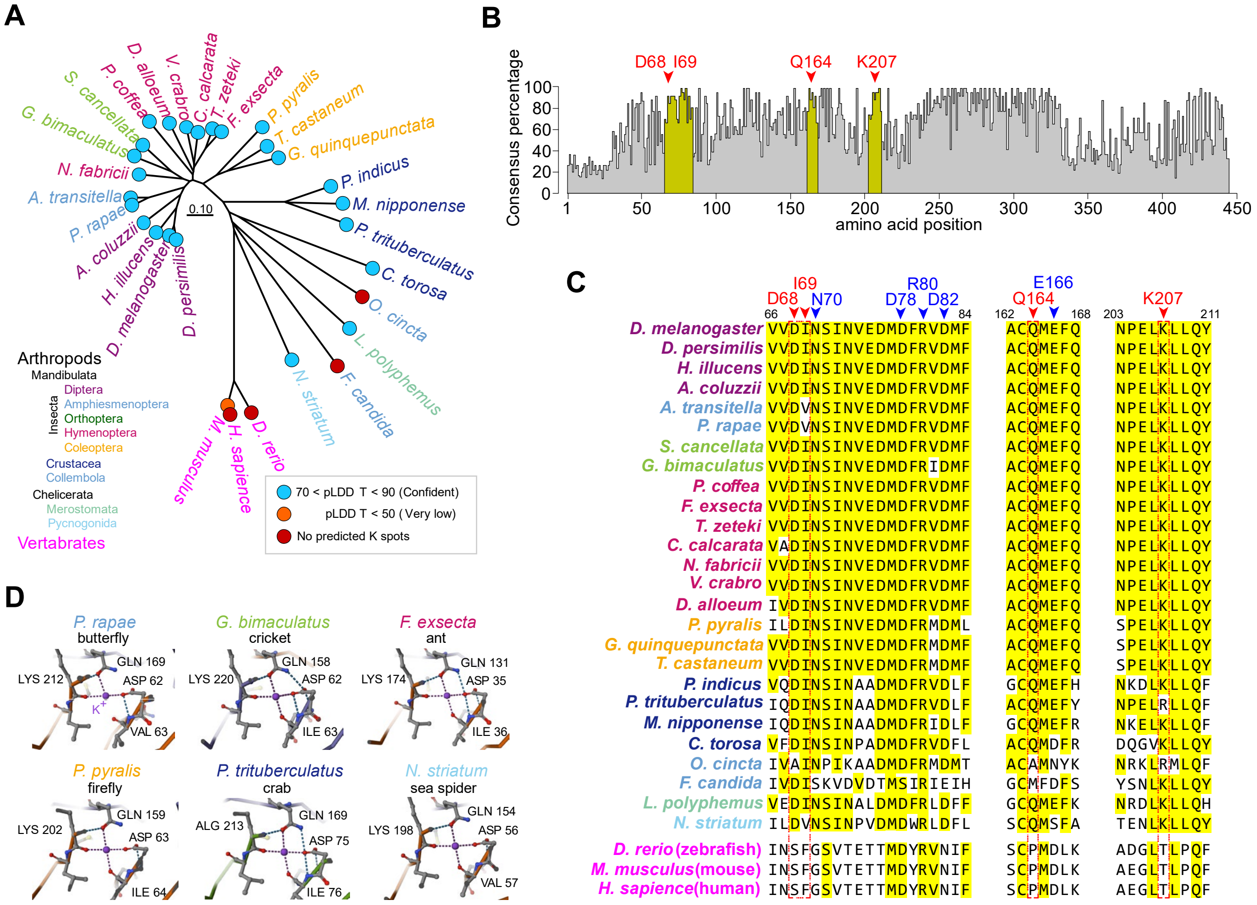


### Fig. S3. Phylogenetic analysis of DmAlka homologs.

(**A**) A phylogenetic tree of the homologous channels of DmAlka in arthropods and vertebrate species. The colored circles at each branch end indicate the pLDDT scores calculated by AF3 for $K^+$. Red color indicates that no $K^+$-spot structures were predicted at the homologous position of DmAlka. (**B**) Amino acid consensus percentage of DmAlka among the arthropod species shown in (A). The gaps between aligned sequences were excluded. The red arrows indicate the amino acid residues that form directly the $K^+$ binding site. (**C**) Sequence alignments of DmAlka and its homologs of arthropods and vertebrate species. The sequence in the region indicated yellow in (B) is shown. Upper numbers indicate the amino acid residues of DmAlka that form directly the $K^+$ binding site (red) or participate in the polar interaction surrounding it (blue). (**D**) Examples of the predicted structures of the $K^+$ binding site in each homolog. $K^+$ are shown in purple.

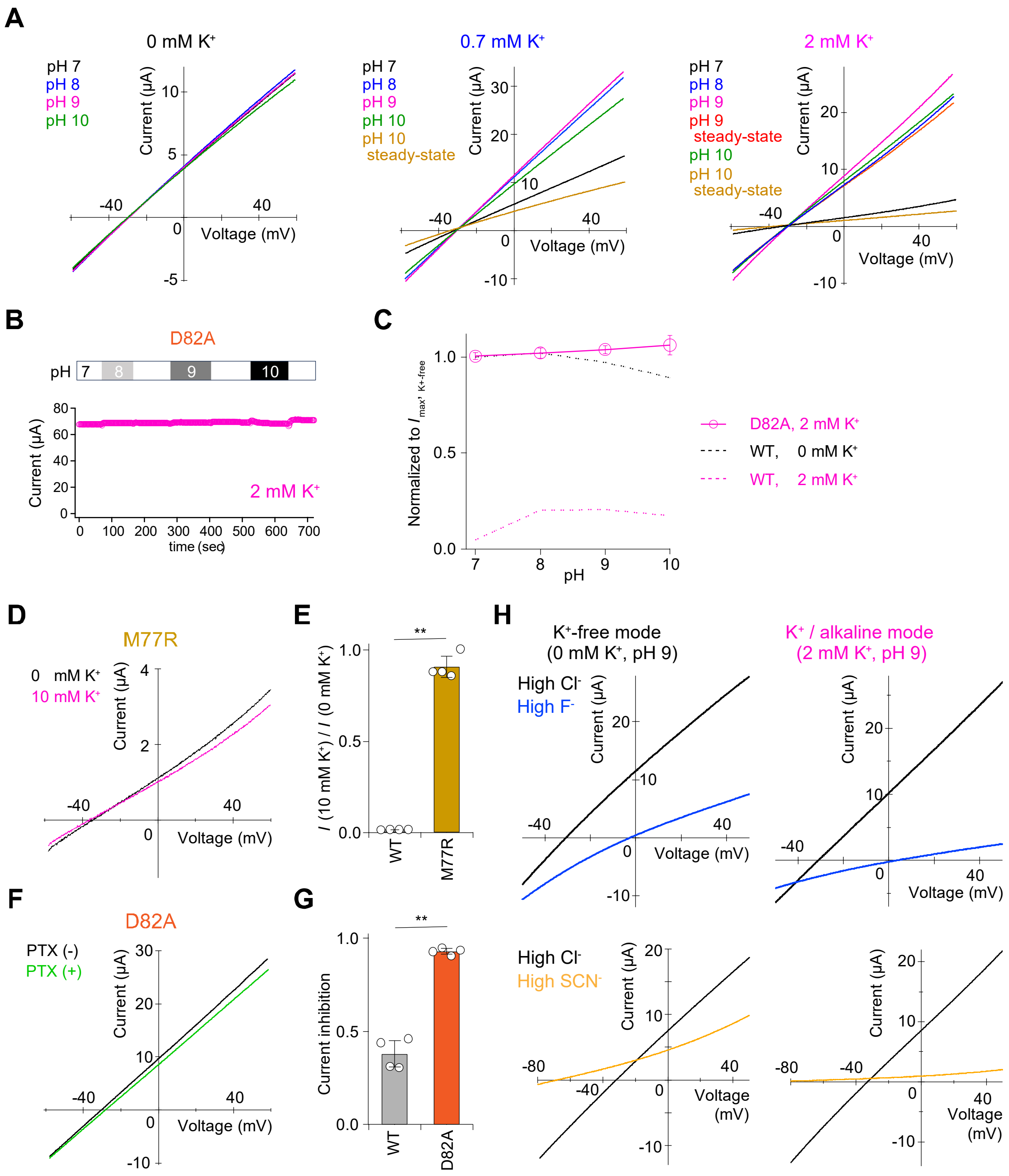
A
0 mM K+
0.7 mM K+
2 mM K+
pH 7
pH 8
pH 9
pH 10
pH 10 steady-state
pH 9 steady-state
Current (μA)
Voltage (mV)
B
D82A
pH 7 8 9 10
Current (μA)
2 mM K+
time (sec)
C
Normalized to I max, K+-free
pH
D82A, 2 mM K+
WT, 0 mM K+
WT, 2 mM K+
D
M77R
0 mM K+
10 mM K+
E
I (10 mM K+) / I (0 mM K+)
WT
M77R
F
D82A
PTX (-)
PTX (+)
G
Current inhibition
WT
D82A
H
K+-free mode (0 mM K+, pH 9)
K+ / alkaline mode (2 mM K+, pH 9)
High Cl-
High F-
High SCN-

**Fig. S4. Extracellular $K^+$-dependent modulation of alkaline sensitivity and pore properties of DmAlka.**

(**A**) Representative current traces recorded under 0-, 0.7-, or 2-mM extracellular $K^+$, at pH 7 (black), pH 8 (blue), pH 9 (magenta), and pH 10 (green). Desensitization was observed under 0.7- or 2-mM extracellular $K^+$, in which peak and steady-state currents (pH 9 in red and pH 10 in brown, respectively) are additionally shown. (**B**) The time-dependent change in the current amplitude at +60 mV in response to alkalization, from pH 7 to pH 8–10, in the D82A mutant in the presence of 2 mM $K^+$. (**C**) The plots of pH-dependence of the D82A currents normalized to those observed in the absence of $K^+$ obtained from (B) (open circle with magenta solid line) (n = 3). The plots of pH-dependence in WT under 0 mM $K^+$ (black dashed line) and 2 mM $K^+$ (magenta dashed line) were obtained from the data in Fig. 3B. (**D**) Representative current traces in the M77R mutant in the absence (black) or presence (magenta) of 10 mM extracellular $K^+$. (**E**) Plot of current reduction by 10 mM $K^+$ in WT and M77R (n = 4). (**F**) Representative current traces of the D82A mutant with (green) or without (black) 1 mM picrotoxin. (**G**) Plot of current inhibition by 1 mM picrotoxin in WT and D82A (n = 4). (**H**) Representative current traces under the bath solutions containing high $Cl^-$ (black), $F^-$ (blue), or $SCN^-$ (brown), composed of each 96 mM sodium anions instead of 96 mM NaCl in ND96. Left and right plots show the data from the $K^+$-free mode (2 mM $NMDG^+$, pH 9) and those from the $K^+$-induced alkaline-activated condition (2 mM $K^+$, pH 9), respectively. Upper panels are for the comparisons between high $Cl^-$ and high $F^-$, and lower panels are for those between high $Cl^-$ and high $SCN^-$.

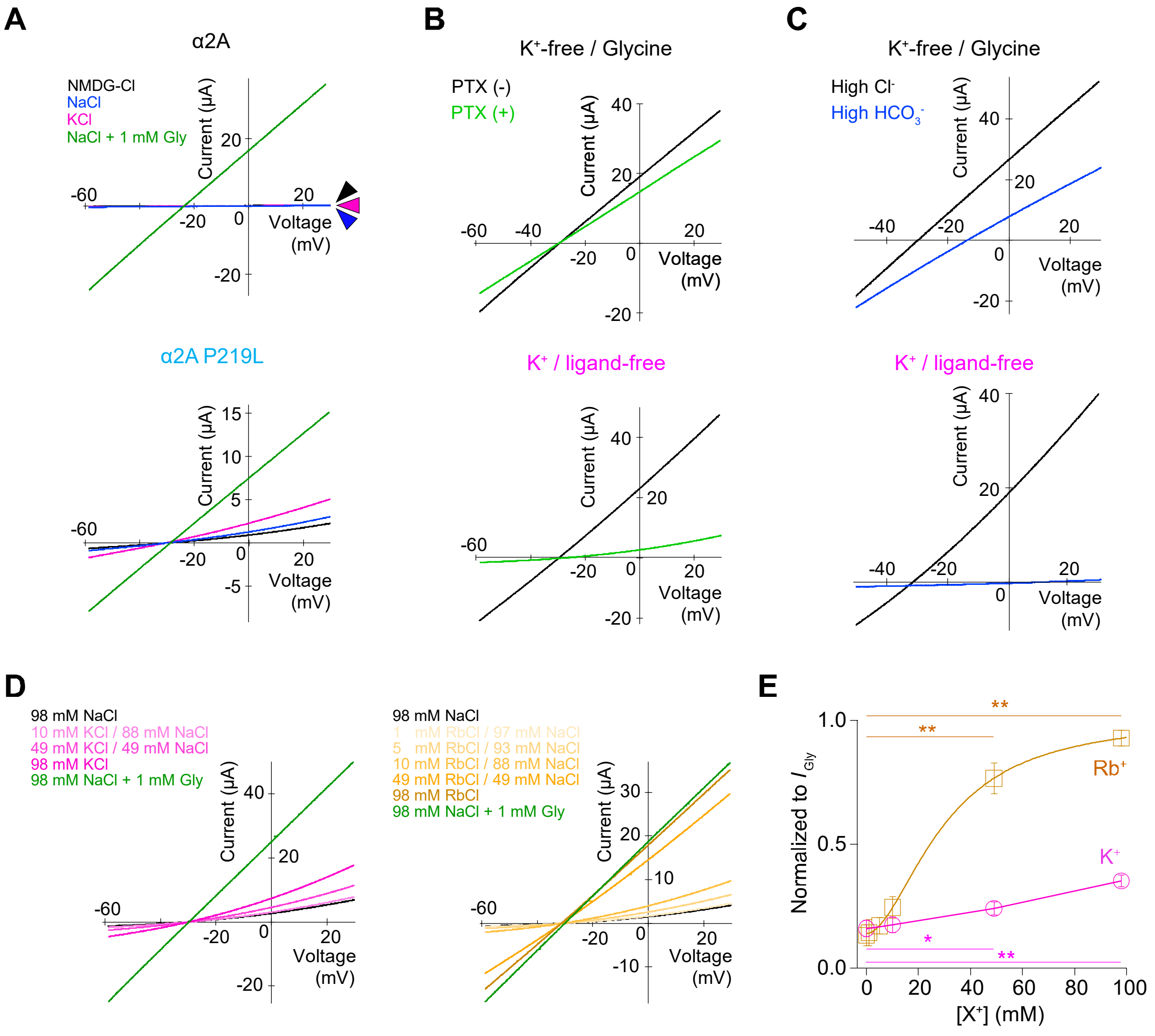


**Fig. S5. Representative current traces of HsGlyRα2B P219L.**

(**A**) Representative current traces in HsGlyRα2A (top) and α2A P219L (bottom) recorded under the basic solutions containing each of 98 mM NMDG-Cl (black), KCl (magenta), NaCl (blue), or NaCl with 1 mM glycine (green). (**B**) Representative current traces of HsGlyRα2B P219L in the presence or absence of 10 μM picrotoxin (PTX), in glycine-induced mode (top) and $K^+$-induced mode (bottom). (**C**) Representative current traces under the bath solution containing high $Cl^-$ (black) or high $HCO_3^-$ (blue), in glycine-induced mode (top) and $K^+$-induced mode (bottom). (**D**) Representative current traces of HsGlyRα2B P219L, showing the dose-dependent changes by $K^+$ (left) and $Rb^+$ (right). The solutions contained 0, 1, 5, 10, 49, or 98 mM KCl (magenta) or RbCl (brown), mixed with NaCl to a final concentration of 98 mM, and 98 mM NaCl with 1 mM glycine (green). (**E**) Plots of dose-response of $Rb^+$ or $K^+$ in HsGlyRα2B P219L. Currents at +30 mV obtained from (D) were used. Currents were normalized to those induced by 1 mM glycine and plotted. The dose response of $Rb^+$ was fitted by Hill equation ($EC_{50} = 29.7 \pm 0.27$ mM), while that of $K^+$ could not be confidently fitted. Statistically significant differences in both $K^+$ and $Rb^+$ were evaluated by comparing the values under 0 mM with those under different concentrations of $K^+$ or $Rb^+$, respectively.

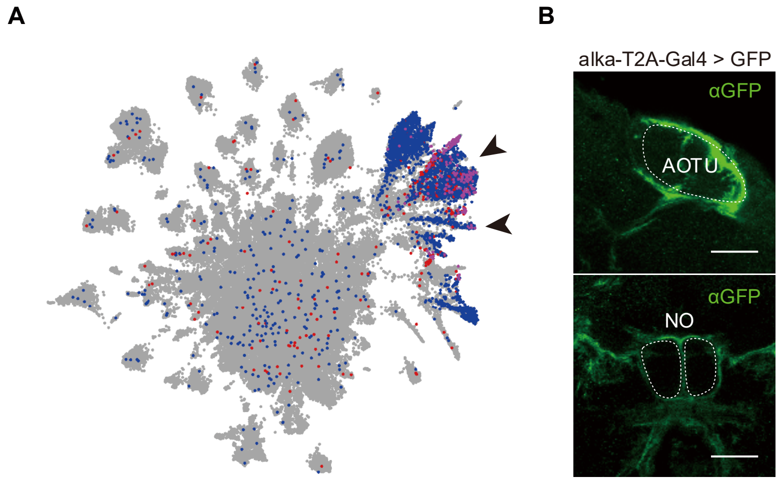


**Fig. S6. Transcriptomic and immunohistochemistry of DmAlka.**

(**A**) Expression profile of alka gene in the adult *Drosophila* brain based on the single-cell transcriptomic data. alka expressing cells are shown in red. Cells that express repo gene, a well-known genetic marker for glial cells, are also depicted in blue. The cells co-expressing both alka and repo are represented in violet. The alka gene is expressed in glial clusters indicated by arrowhead. Images sourced from the database SCope (*6*) (http://scope.aertslab.org) are shown. (**B**) The alka positive cells were visualized by expressing GFP from alka-T2A-Gal4 driver (*7*). The alka-T2A-Gal4 is a bicistronic driver in which the self-cleaving T2A peptide and Gal4 are knocked into immediately before the stop codon of the alka gene, yielding both the native DmAlka protein and Gal4 simultaneously. Thus, Gal4-induced GFP signals reflect DmAlka translation. GFP signals exhibited the typical morphology of glial cells that surround neuropiles, including the anterior optic tubercle (AOTU) and the nodulus (NO) in the brain. Scale bar, 20 μm.

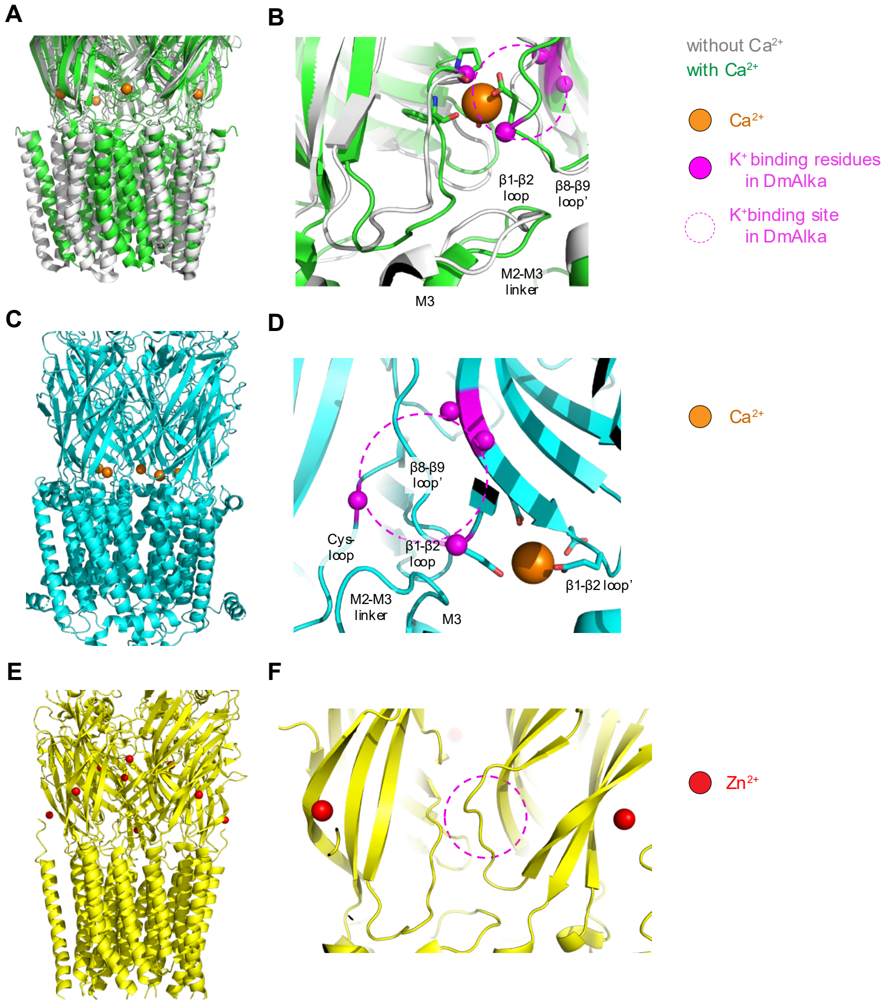


**Fig. S7. Structural comparisons with other pLGIC structures.**

(**A**) An overlay of the $Ca^{2+}$-free (white) and $Ca^{2+}$-bound (green) structures of DeCLIC, shown as a cartoon representation (PDB: 6V4S and 6V4A). Brown spheres represent bound $Ca^{2+}$. (**B**) A close-up view of the $Ca^{2+}$-binding site in DeCLIC. Magenta spheres indicate the positions of the $C_\alpha$ atoms corresponding to those forming the $K^+$-binding sites in DmAlka. The residues in DeCLIC that directly coordinate $Ca^{2+}$ are shown as stick models. The region for the $K^+$-binding site in DmAlka is roughly shown as magenta circles with dotted line. (**C**) A structure of human nicotinic acetylcholine receptor (AChR) α7 (PDB: 7KOX), colored in aqua. Brown spheres represent bound $Ca^{2+}$. (**D**) A close-up view of the $Ca^{2+}$-binding site in nAChR, shown as similar to (B). (**E**) A structure of human GlyR α3 (PDB: 9BWJ), colored in yellow. Red spheres represent bound $Zn^{2+}$. (**F**) A close-up view of the $Zn^{2+}$-binding site in GlyR α3, shown as similar to (B).

**Table S1. Anion selectivity in DmAlka and HsGlyRα2B P219L.**

Selectivity of each anion relative to $Cl^-$ calculated for DmAlka and HsGlyRα2B P219L by exchanging the bath solution containing high $Cl^-$ and high anions. For DmAlka, comparisons were made in the $P_{Cl-}$ / $P_x$ values between the $K^+$-induced and alkaline-activated mode (2 mM $K^+$ / pH 9) and the $K^+$-free mode (2 mM $NMDG^+$ / pH 9). For HsGlyRα2B P219L, the $P_{Cl-}$ / $P_{HCO3-}$ values were compared between the conditions recorded under 98 mM $K^+$ and under 98 mM NaCl with 0.1 mM glycine. For experimental details, see Materials and Methods.

**$P_{x^-}$ / $P_{Cl^-}$**

| | Average | | S.D. | n | Average | | S.D. | n | significance |
|---|---|---|---|---|---|---|---|---|---|
| ***DmAlka*** | | | | | | | | | |
| | $K^+$ (+) | | | | $K^+$ (-) | | | | |
| $HCO_3^-$ / $Cl^-$ | 0.253 | ± | 0.018 | 6 | 0.319 | ± | 0.023 | 6 | ** (0.00046) |
| $F^-$ / $Cl^-$ | 0.250 | ± | 0.021 | 4 | 0.346 | ± | 0.065 | 4 | n. s. (0.051) |
| $I^-$ / $Cl^-$ | 1.49 | ± | 0.11 | 4 | 1.32 | ± | 0.021 | 4 | n. s. (0.053) |
| $SCN^-$ / $Cl^-$ | 9.74 | ± | 2.3 | 4 | 3.56 | ± | 0.44 | 4 | ** (0.0036) |
| ***HsGlyR α2B P219L*** | | | | | | | | | |
| | $K^+$-mode | | | | Glycine-mode | | | | |
| $HCO_3^-$ / $Cl^-$ | 0.193 | ± | 0.034 | 5 | 0.352 | ± | 0.089 | 4 | * (0.015) |

### References


1. T. Shimomura, Y. Kubo, Phosphoinositides modulate the voltage dependence of two-pore channel 3. *J. Gen. Physiol.* **151**, 986–1006 (2019).

2. J. Abramson, J. Adler, J. Dunger, R. Evans, T. Green, A. Pritzel, O. Ronneberger, L. Willmore, A. J. Ballard, J. Bambrick, S. W. Bodenstein, D. A. Evans, C.-C. Hung, M. O'Neill, D. Reiman, K. Tunyasuvunakool, Z. Wu, A. Žemgulytė, E. Arvaniti, C. Beattie, O. Bertolli, A. Bridgland, A. Cherepanov, M. Congreve, A. I. Cowen-Rivers, A. Cowie, M. Figurnov, F. B. Fuchs, H. Gladman, R. Jain, Y. A. Khan, C. M. R. Low, K. Perlin, A. Potapenko, P. Savy, S. Singh, A. Stecula, A. Thillaisundaram, C. Tong, S. Yakneen, E. D. Zhong, M. Zielinski, A. Žídek, V. Bapst, P. Kohli, M. Jaderberg, D. Hassabis, J. M. Jumper, Accurate structure prediction of biomolecular interactions with AlphaFold 3. *Nature* **630**, 493–500 (2024).

3. T. J. Dolinsky, J. E. Nielsen, J. A. McCammon, N. A. Baker, PDB2PQR: an automated pipeline for the setup of Poisson-Boltzmann electrostatics calculations. *Nucleic Acids Res.* **32**, W665-7 (2004).

4. R. Andrey, N. Masatoshi, A simple method for estimating and testing minimum-evolution trees. *Mol. Biol. Evol.*, doi: 10.1093/oxfordjournals.molbev.a040771 (1992).

5. S. Kumar, G. Stecher, M. Suleski, M. Sanderford, S. Sharma, K. Tamura, MEGA12: Molecular evolutionary genetic analysis version 12 for adaptive and green computing. *Mol. Biol. Evol.* **41** (2024).

6. K. Davie, J. Janssens, D. Koldere, M. De Waegeneer, U. Pech, Ł. Kreft, S. Aibar, S. Makhzami, V. Christiaens, C. Bravo González-Blas, S. Poovathingal, G. Hulselmans, K. I. Spanier, T. Moerman, B. Vanspauwen, S. Geurs, T. Voet, J. Lammertyn, B. Thienpont, S. Liu, N. Konstantinides, M. Fiers, P. Verstreken, S. Aerts, A single-cell transcriptome atlas of the aging Drosophila brain. *Cell* **174**, 982-998.e20 (2018).

7. S. Kondo, T. Takahashi, N. Yamagata, Y. Imanishi, H. Katow, S. Hiramatsu, K. Lynn, A. Abe, A. Kumaraswamy, H. Tanimoto, Neurochemical organization of the Drosophila brain visualized by endogenously tagged neurotransmitter receptors. *Cell Rep.* **30**, 284-297.e5 (2020).
